# Methomyl Injury to Carbon Exchange Rates and Related Processes in Cotton

**DOI:** 10.1101/121798

**Authors:** C.E. Salem, J.T. Cothren, C.R. Benedict

## Abstract

Some insecticides have the potential to cause varying levels of phytotoxicity. This study examined 1) the time courses of photosynthetic injury in cotton (Gossypium hirsutisms L.) leaves treated with methomyl [S-methyl-N-[(methyl carbamoyl)oxy]-thioacetimidate] and 2) the relationships between carbon exchange rate (CER), stomatal conductance, the chlorophyll fluorescence parameter FX/FP, and ribulose-1,5-bisphosphate carboxylase/ oxygenase (rubisco) activity. Plots were sprayed with either 0 or 0.84 kg methomyl·ha-1 when cotton was in mid-reproductive growth. Starting on the day of spraying, CER, stomatal conductance, FX/FP, and rubisco activity were measured daily for five consecutive days [4, 28, 52, 76 and 100 hours after spraying (HAS)]. In methomyl-treated leaves, CER decreased within hours after spraying, reached their lowest point at 28 HAS in experiment I and 76 HAS in experiment II, then recovered near-control levels by 100 HAS. At their lowest points, CER of methomyl-treated leaves decreased from 20 to 50% compared to controls. Stomatal conductance, FX/FP, and rubisco activity followed similar patterns to CER. Stomatal conductance was more closely related to CER than were FX/FP and rubisco activity. Chlorophyll fluorescence recovered more quickly than did CER. Rubisco activity did not decrease till after CER. From the parameters measured in this study, stomatal conductance appeared to be the major factor influencing methomyl-induced changes in CER, although all three parameters may be involved in the process of CER change.

**Abbreviations:**

CER: carbon exchange rate
rubisco: ribulose-1,5-bisphosphate carboxylase/ oxygenase
DAT: days after treatment
HAS: hours after spraying

---

Some insecticides, despite their beneficial effects on insect control, have harmful side effects on the crops. One harmful side effect that may occur is injury to the photosynthetic system, which may significantly reduce crop yields (Haile et al., 1999). Wood and Payne (1986) reported that the carbon exchange rates (CER) of mature pecan [*Carya illionensis (Wang) K. Koch*] were decreased by a single spray at recommended rates for eight out of nine insecticides examined. In their study, CER decreased approximately 20% by one day after treatment (1 DAT), began to recover by 8 DAT, and showed no significant differences by 26 DAT. They also found that wettable powder (WP) formulations generally caused less injury than emulsifiable compound (EC) formulations, but the difference was significant only for methomyl EC and WP at 1 DAT. Wood and Payne (1986) concluded, however, that stomatal conductance was generally unaffected by any insecticide treatment. Youngman (1990) found that carbon exchange rates (CER) and stomatal conductance of cotton were not significantly affected by treatment with methyl parathion, propargite, chlordimeform, permethrin, or methomyl. However, methyl parathion and permethrin decreased mesophyll conductance at 3 DAT, and recovery was shown later.

Chlorophyll fluorescence can be used to monitor stress on the photosynthetic system (Maxwell and Johnson, 2000). A chlorophyll fluorescence transient generally consists of a fast phase followed by a slow phase. For reviews of chlorophyll fluorescence parameters and their significance, see Krause and Weis (1991), Lichtenthaler (1988), Maxwell and Johnson (2000), and Papageorgiou (1975). Fluorescence parameters from fast fluorescence have often been used as estimates of photosynthetic strength (Maxwell and Johnson, 2000). The most commonly used parameters of chlorophyll fluorescence are the ratio of variable fluorescence (F_V_) from dark-adapted leaves to maximal fluorescence (F_P_), F_V_/F_P_, which is a measure of the maximum quantum yield of photosystem II (PSII), and the ratio of variable fluorescence from dark-adapted leaf to minimal fluorescence (F_O_), F_V_/F_O_. In studies of penetration of photosynthetically active herbicides into leaf tissue, all three of these parameters (F_V_/F_P_, F_V_/F_O_, and F_X_/F_P_) decreased with increasing photosynthetic injury, and the ratio of variable fluorescence (F_X_) from light-adapted leaf to maximal fluorescence, F_X_/F_P_, was determined to be the best indicator of photosynthetic strength (Habash et al., 1985).

Herbicidal photosynthesis inhibitors (e.g., diuron) greatly increased minimal fluorescence of dark- and light-adapted leaves (F_O_ and F_I_), thus decreasing F_V_ and F_X_, even though F_P_ often rises. With total positioning of the system, F_P_ rises swiftly to a maximum value and keeps consistent; the fast or slow fluorescence transient does not occur (Habash et al., 1985).

Stresses that affect efficiency of PS II lead to a characteristic decrease in F_V_/F_P_ (Krause and Weis, 1991). This decrease is usually a result of decreased F_V_ and very often of increased F_O_ (Lichtenthaler, 1988). The factors causing these and related changes in fast fluorescence induction include contact herbicides (Szigeti et al., 1988), Mn deficiency (Simpson and Robinson, 1984), chilling injury (Bolhar-Nordenkampf and Lechner, 1988; Greaves and Wilson, 1987; Larcher and Neuner, 1989), and moisture stress (Ben et al., 1987; Genty et al., 1987; Ogren, 1990; Chen et al., 2012a, 2012b; Wen et al., 2013).

Effects of methomyl on cotton include increased foliage reddening, premature defoliation, chlorophyll degradation, ethylene evolution, and electrolyte leakage. In a field study conducted by Lincoln and Dean (1976), high methomyl rate treatment resulted in foliage reddening and moderate late-season defoliation. Methomyl increased anthocyanin and tannin content, and decreased chlorophyll content in mature leaves rather than expanding leaves (Parrott et al., 1983). Ethylene evolution increased significantly in cotton leaf within 9 h (1 h + 8 h incubation, as leaf discs) and remained higher than the ethylene level of controls till 144 h after treatment (Guthrie and Cothren, 1989). Guthrie and Cothren (1989) further concluded that since ethylene evolution was stimulated before electrolytic leakage, the electrolytic leakage may be attributed to altered membrane permeability associated with enhanced ethylene production. Most of the above effects are often associated with senescence, suggesting that methomyl may hasten senescence. Despite the various effects of methomyl on cotton leaves, it usually does not decrease yields significantly (Durant, 1977; Lincoln and Dean, 1976). Methomyl toxicity from the dust formation is considerably less than from aqueous formulations due to decreased absorption (Bull, 1974).

In addition, in our preliminary studies, methomyl treatment temporarily injured the photosynthetic system of cotton leaves (Salem et al., 1990), as indicated by decreases in the chlorophyll fluorescence parameter F_X_/F_P_. This injury to chlorophyll fluorescence led us to examine carbon exchange rates and other related processes. The objectives in this experiment were to establish the time courses of photosynthetic injury and leaf reddening in methomyl-treated cotton leaves, and to examine the relationships between carbon exchange rate, stomatal conductance, chlorophyll fluorescence, and ribulose-1,5-bisphosphate carboxylase activity.

## MATERIALS AND METHODS

Two field experiments were conducted at the Texas A&M Research Farm in Burleson County, Texas in 1991. In experiment I, cotton cultivar Stoneville 453 was planted on April 11 in a Weswood silt loam (*Fluventic Ustochrept*). In experiment II, cotton cultivar Deltapine 50 was planted on April 30 on a Ships clay (*Udic Chromustert*). Standard agronomic practices for the area were followed. Plots were sprayed with either 0 or 0.84 kg·ha^-1^ methomyl (control and methomyl, respectively) before 1000 h CDT on July 8 and July 29 in experiments I and II, respectively. Both experiments were designed as randomized complete blocks with two treatments (control or methomyl) and four replications.

The youngest fully expanded leaf at time of spraying was tagged for subsequent readings on five plants in each plot. Carbon exchange rate (CER), stomatal conductance (COND), and ribulose-1,5-bisphosphate carboxylase activity (Rubisco) were repeatedly measured at approximately 4, 28, 52, 76, and 100 hours after spraying (HAS). For rubisco, the leaves were sampled at the times noted above, and the measurements were taken in the laboratory (as explained below). The chlorophyll fluorescence parameter F_X_/F_P_ was measured at 20, 44, 68, and 92 HAS; the timing of the chlorophyll fluorescence readings was due to the necessity of pre-dawn measurements.

Chlorophyll fluorescence was measured using an SF-30 Fluorometer (Richard Brancker Ltd., Ottawa, Canada). This instrument, which uses a small probe, cannot provide whole-leaf readings. Hence, the chlorophyll fluorescence measurements were taken on non-spotted portions (i.e. showing no visible injury) of the tagged leaves. The probe was positioned in the top-half of a dark-adapted leaf between the midrib and the margin. Light of 670 nm wavelength (approximately 8 W·m^-2^ in intensity) was beamed onto a 20-mm^2^ section of the leaf. The fluorometer monitored the intensity of the returning light at 10-ms intervals for 10 s, then stored the resulting trace (Anonymous, 1986). Based on this trace, the ratio F_X_/F_P_ was calculated as described by Habash et al. (1985).

Stomatal conductance and CER were measured between 1200 and 1400 h CDT with a Li-Cor 6200 Portable Photosynthesis System (Li-Cor, Inc.; Lincoln, NE) on overcast days. The readings were not taken until the sun was unobscured by clouds for at least 5 min. A 4-L assimilation chamber was clamped onto the leaf until CO_2_ concentrations decreased at least 35 μL·L^-1^ from ambient. Both stomatal conductance and CER were calculated by the equations given in the LI-6200 Technical Reference (Anonymous, 1987).

After CER and stomatal conductance were determined, the leaf was transported on ice to the laboratory. Then, the leaf was video-imaged if visible leaf injury had occurred. The leaf fresh weight was weighed, and its area was scanned using a leaf area meter. The rubisco activity of the leaf was also assayed. The leaf spotting in percentage of the leaf area was determined by image capture analysis on the video-images of the leaves (Image Capture Analysis System; Digital Image Acquisition Systems, Inc., Englewood, CO). Percent spotting included not only reddening, but also chlorosis or necrosis. Attempts to separate the three types of visible injury were not successful because brown and red were not distinguishable on the video-images. In almost all cases, visibly injured areas in methomyl-treated leaves were red. These reddened areas were deemed to be injured rather than only reddened, because reddened areas produced chlorophyll fluorescence values similar to those of necrotic tissue (data not shown).

After the leaves were video-imaged and weighed, they were used to determine rubisco activity. Due to time limitation only a portion of the leaf was used in the assay. Because the leaf reddening was usually not symmetrical, the whole leaf was sliced into small pieces and well mixed, then a randomly taken 1.25 g portion was used for the assay. Rubisco activity was assayed according to Wong and Benedict (1983) with the following modifications. Trition-X 100 [octylphenoxy-poly(ethoxy)ethanol] was added to the grinding medium at 1.0% (v:v) to solubilize the enzyme. After the initial centrifugation (at 27000 g for 30 min), pellets were resuspended in 5 ml grinding medium and centrifuged twice more (at 27000 g for 10 min).

Analysis using the full statistical model, although useful, was inadequate for these experiments for the following reasons. First, owing to the injury and subsequent recovery of methomyl-treated plants, the Methomyl × Hour interactions are of more importance than the Methomyl or Hour main effect *per se*. Second, and corollary to the first reason, interactions not including both methomyl and hour (e.g. Methomyl × Experiment and Hour × Experiment) were usually either not of interest or difficult to explain. Third, the relationships between dependent variables (CER, stomatal conductance, chlorophyll fluorescence, and rubisco activity) are of great interests. Normally, only the significant interactions from the full model would be further analyzed. However, since some interactions were significant for one dependent variable, but not for others (e.g. Experiment × Methomyl × Hour was significant for CER, but not for stomatal conductance, chlorophyll fluorescence, or rubisco), breaking out only the significant interactions would not allow full comparison of all dependent variables. To address the preceding problems, the Methomyl×Hour interaction (for each dependent variable) was graphed for each experiment. Furthermore, at each hour, a mean comparison between control and methomyl was made. The full statistical model is presented but generally not discussed unless a particular main effect or interaction was of importance. All statistical analyses were performed in SAS 9.2 (SAS Institute, Cary, NC).

## RESULTS

### Leaf Spotting

Methomyl-induced spotting in leaves typically appears as interveinal and marginal reddening. Spotting did not occur until 52 HAS in experiment II, but occurred by 28 HAS in experiment I (Table 1). In both experiments, spotting peaked at 52 HAS. Spotting was relatively mild, averaging approximately 5% of the leaf area.

**Table 1.** Effects of methomyl (MET) on spotting in cotton leaves at various hours after spraying (HAS). Exp I and II signify experiment I and II, respectively.

| **Exp I** |  | **SPOTTING** |  |  |  |  |
| --- | --- | --- | --- | --- | --- | --- |
| **MET** |  | 4<br>HAS | 28<br>HAS | 52<br>HAS | 76<br>HAS | 100<br>HAS |
| kg·ha<sup>-1</sup> |  |  |  | % |  |  |
| 0.00 |  | . | 1.0 | 1.9 | 1.2 | 0.6 |
| 0.84 |  | . | 4.1 | 8.6 | 4.0 | 4.5 |
| **Significance** |  |  | ** | + | * | + |

| **Exp II** |  | **SPOTTING** |  |  |  |  |
| --- | --- | --- | --- | --- | --- | --- |
| **MET** |  | 4<br>HAS | 28<br>HAS | 52<br>HAS | 76<br>HAS | 100<br>HAS |
| kg·ha<sup>-1</sup> |  |  |  | % |  |  |
| 0.00 |  | . | . | 1.9 | 1.9 | 1.9 |
| 0.84 |  | . | . | 8.6 | 5.1 | 4.0 |
| **Significance** |  |  |  | + | NS | NS |

\*\*, \*, +, NS represent $p < 0.01$ , $p < 0.05$ , $p < 0.1$ , and $p > 0.1$ , respectively.

### Carbon Exchange Rates (CER)

In experiment I, CER of control plants declined gradually from 24.1 μmol CO_2_ m^-2^·s^-1^ at 4 HAS to 20.3 μmol CO_2_ m^-2^·s^-1^ at 100 HAS (Fig. 1A). These values are within reported CER ranges in cotton (Peng and Krieg, 1991; Wullschleger and Oosterhuis, 1990; Youngman et al., 1990). In addition, these values are quite similar to the values reported in previous studies (Bauer, 1988) in the same geographical area. For methomyl-treated plants, CER had already decreased significantly compared to the control at 4 HAS. The lowest reading ascertained for methomyl-treated plants occurred at 28 HAS and represented a decrease of nearly 50% compared to control plants. From that point on, CER recovery started, and by 76 HAS the CER of methomyl-treated and control plants did not differ significantly.

**Figure 1.**
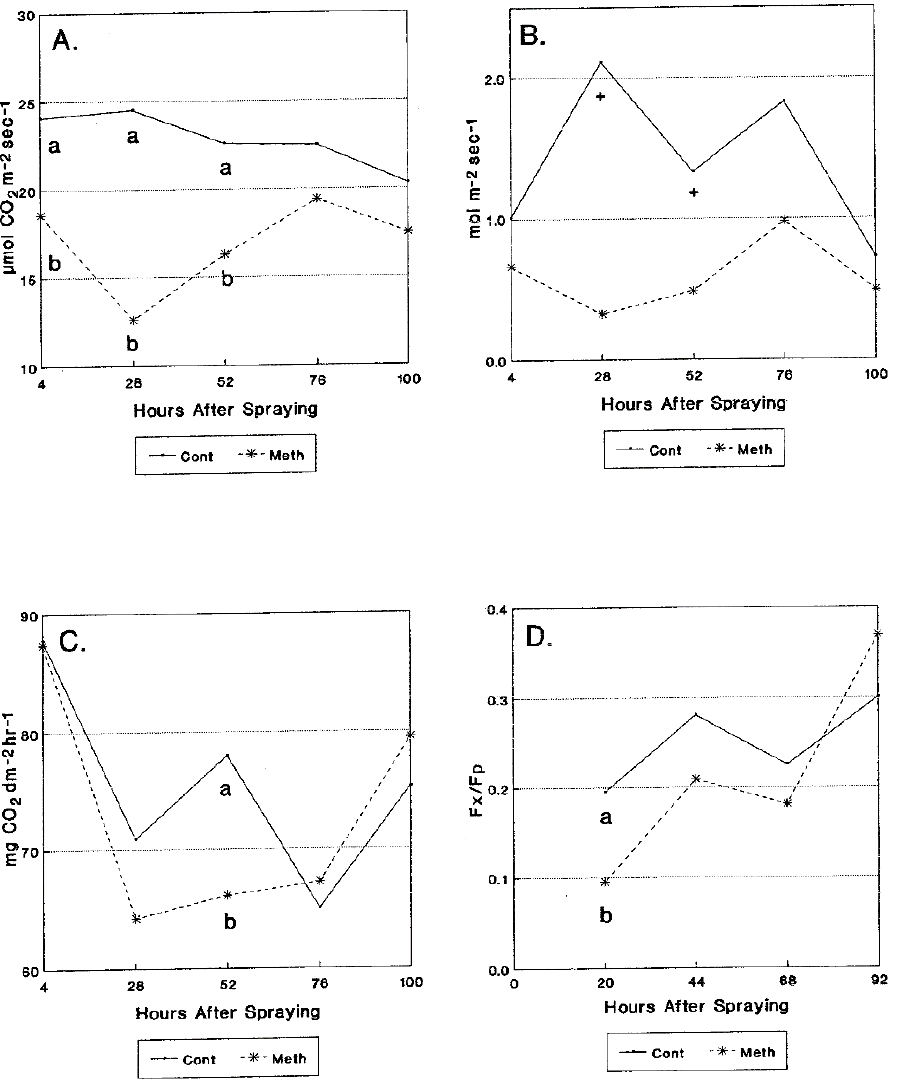
The effects over time of methomyl on (A) carbon exchange rates, (B) stomatal conductance, (C) rubisco activity, and (D) the chlorophyll fluorescence parameter F_x_/F_p_ of the youngest fully expanded leaf of cotton cultivar Stoneville 43. Cont and Meth = 0.00 and 0.84 kg methomyl ha^-1^, respectively. Values at a given hour with the same letter do not differ significantly at p < 0.05 according to Duncan’s Multiple Range Test. + signifies differences at p < 0.1.

In experiment II, CER of control plants increased gradually from 24.7 μmol CO_2_ m^-2^· s^-1^ at 4 HAS to 26.4 μmol CO_2_ m^-2^·s^-1^ at 100 HAS (Fig. 2A). Again, these values are within the reported ranges for cotton (Peng and Krieg, 1991; Wullschleger and Oosterhuis, 1990; Youngman et al., 1990). The CER of methomyl-treated plants were lower (p < 0.1) than those of control plants at 4 and 28 HAS, and significantly lower (p < 0.05) at 52 and 76 HAS. At 100 HAS, the difference in CER between control and methomyl-treated plants became non-significant. The lowest CER for methomyl-treated plants occurred at 76 HAS. At this point, CER decreased approximately 27% in methomyl-treated plants compared to controls.

**Figure 2.**
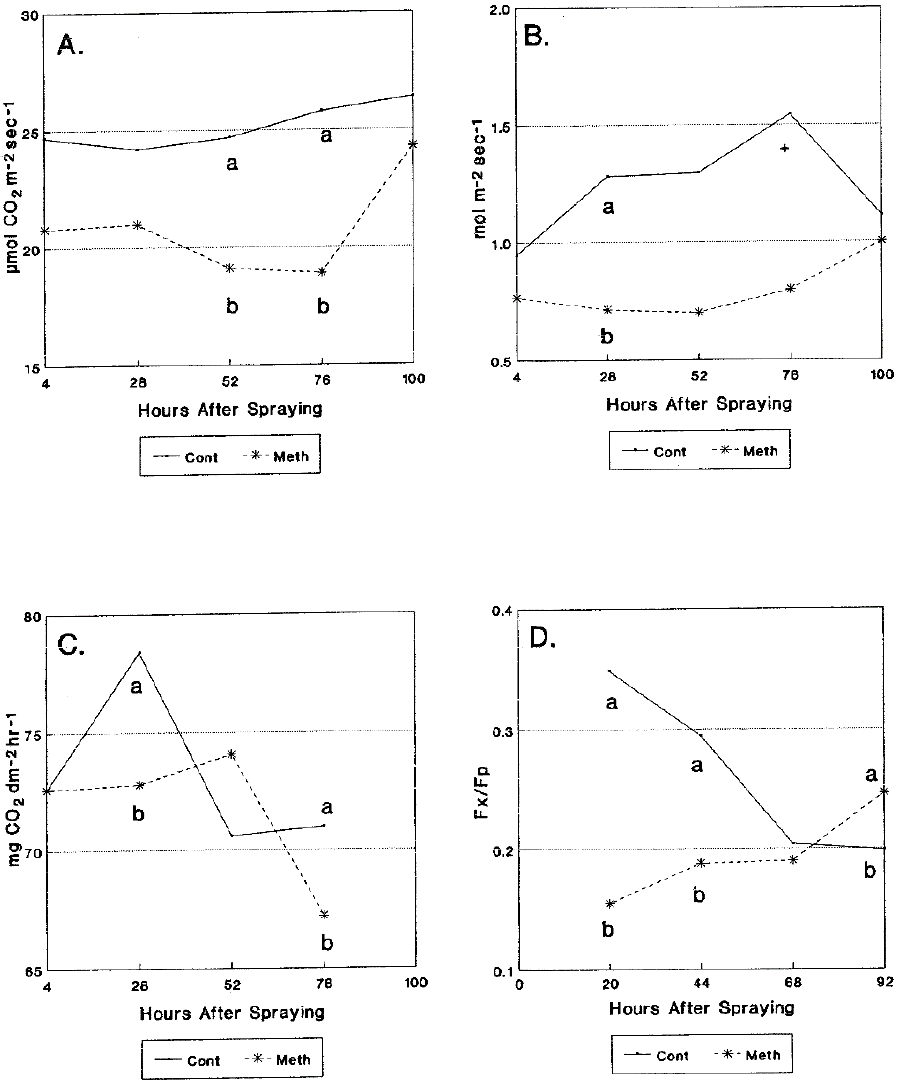
The effects over time of methomyl on (A) carbon exchange rates, (B) stomatal conductance, (C) rubisco activity, and (D) the chlorophyll fluorescence parameter F_x_/F_p_ of the youngest fully expanded leaf of cotton cultivar Deltapine 50. Cont and Meth = 0.00 and 0.84 kg methomyl ha-1, respectively. Values at a given hour with the same letter do not differ significantly at p < 0.05 according to Duncan’s Multiple Range Test. + signifies differences at p < 0.1.

### Stomatal Conductance

In experiment I, stomatal conductance of control plants varied from 2.1 to 0.7 mol·m^-2^·s^-1^ (Fig. 1B). These values are somewhat higher than previously reported values (Peng and Krieg, 1991; Wullschleger and Oosterhuis, 1990; Youngman et al., 1990), but were similar to the values reported by Bauer (1988). In methomyl-treated plants, stomatal conductance followed a similar pattern as CER: the lowest reading (0.33 mol·m^-2^·s^-1^) occurred at 28 HAS, and by 76 HAS the stomatal conductance of methomyl-treated plants did not differ significantly from that of control plants. At 28 and 52 HAS (when methomyl-treated and control plants differed at p < 0.1), stomatal conductance decreased 85 and 64%, respectively, in methomyl-treated plants compared to controls.

In experiment II, stomatal conductance of control plants varied from 0.95 to 1.55 mol·m^-2^·s^-1^ (Fig. 2B). At 28 and 76 HAS, stomatal conductance of methomyl-treated plants decreased 44 and 48%, respectively, compared with controls. At 100 HAS, stomatal conductance values of control and methomyl-treated plants were not significantly different.

### Rubisco Activity

In experiment I, rubisco activity of control plants ranged from 67 to 88 mg CO_2_ dm^-2^·hr^-1^ (Fig. 1C). These values were slightly higher than expected. Mahan et al. (1984), working with diploid species of cotton, reported rubisco activities from 4.35 to 76 mg CO_2_ dm^-2^·hr^-1^. Rubisco activity in methomyl-treated plants first declined then recovered. The decrease was significant only at 52 HAS. At this point, rubisco activity declined approximately 15% in methomyl-treated plants compared to controls. After this point, rubisco activities of methomyl-treated plants were numerically, but not significantly, greater than that of controls.

In experiment II, the 100 HAS readings were not obtained (Fig. 2C). At 28 and 76 HAS, rubisco activity decreased by approximately 7% and 5%, respectively, in methomyl-treated plants compared with controls. At 52 HAS, rubisco activity was higher in the methomyl-treated plants than in the controls, but the difference was not significant. The decreased rubisco activity apparent in both experiments is most readily explained by injury to the photosynthetic system. However, since the decrease is temporary, the injury to the photosynthetic system would also appear to be temporary.

Despite the individual differences found in each experiment at certain hours, neither the full model (Table 2) nor either of the “by experiment” models (Table 3) was able to detect any effect of methomyl on rubisco activity.

**Table 2.** Analysis of variance for the effects of experiment (EXP), methomyl rate (MET), and hour (HR) on carbon exchange rates (CER), stomatal conductance (COND), chlorophyll fluorescence (F_x_/F_p_), and ribulose-1,5-bisphosphate carboxylase activity (Rubisco).

| **FACTOR** | **CER** |  | **COND** |  | **F<sub>x</sub>/F<sub>p</sub>** |  | **Rubisco** |  |
| --- | --- | --- | --- | --- | --- | --- | --- | --- |
|  | **df** | **p-value** | **df** | **p-value** | **df** | **p-value** | **df** | **p-value** |
| **REP** | 3 | 0.1618 | 3 | 0.1986 | 3 | 0.3619 | 3 | 0.9095 |
| **EXP (E)** | 1 | 0.0225 * | 1 | 0.7747 | 1 | 0.8735 | 1 | 0.7932 |
| **Error A** | 3 | - | 3 | - | 3 | - | 3 | - |
| **MET (M)** | 1 | 0.0003 ** | 1 | 0.0001 ** | 1 | 0.1083 | 1 | 0.9236 |
| **M×E** | 1 | 0.3025 | 1 | 0.0425 * | 1 | 0.2972 | 1 | 0.5978 |
| **Error B** | 6 | - | 6 | - | 6 | - | 6 | - |
| **HR (H)** | 4 | 0.0514 + | 4 | 0.3007 | 3 | 0.0123 * | 3 | 0.0005 ** |
| **H×E** | 4 | 0.0014 ** | 4 | 0.5687 | 3 | 0.0030 ** | 3 | 0.0010 ** |
| **H×M** | 4 | 0.0077 ** | 4 | 0.2236 | 3 | 0.0011 ** | 3 | 0.6573 |
| **H×M×E** | 4 | 0.0006 ** | 4 | 0.7409 | 3 | 0.8957 | 3 | 0.2601 |
| **Error C** | 48 | - | 48 | - | 36 | - | 36 | - |

\*\*, \*, +, NS represent $p < 0.01$ , $p < 0.05$ , $p < 0.1$ , and $p > 0.1$ , respectively.

**Table 3.** Analysis of variance for the effects of methomyl rate (MET) and hour (HR) on ribulose-1,5-bisphosphate carboxylase activity (Rubisco). Separate analyses of variance for experiments I and II (Exp I, Exp II).

| FACTOR | Exp I |  |  | Exp II |  |
| --- | --- | --- | --- | --- | --- |
|  | df | p-value |  | df | p-value |
| **REP** | 3 | 0.4322 |  | 3 | 0.4776 |
| **EXP (E)** | 1 | 0.5363 |  | 1 | 0.5075 |
| **Error A** | 3 | - |  | 3 | - |
| **MET (M)** | 4 | 0.0012 | ** | 3 | 0.0277 * |
| **M×E** | 4 | 0.4675 |  | 3 | 0.1190 |
| **Error B** | 24 | - |  | 18 | - |

\*\*, \*, +, NS represent $p < 0.01$ , $p < 0.05$ , $p < 0.1$ , and $p > 0.1$ , respectively.

### Chlorophyll Fluorescence

In experiment I, the chlorophyll fluorescence parameter F_X_/F_P_ decreased 51% at 20 HAS in methomyl-treated plants (Fig. 1D). As mentioned above, decreases in F_X_/F_P_ signify decreases in photosynthetic efficiency. The control and methomyl-treated plants did not differ significantly at any other time; however, at 92 HAS, chlorophyll fluorescence was numerically higher in methomyl-treated plants than in controls.

In experiment II, chlorophyll fluorescence of control plants decreased from 20 to 92 HAS. At 20 and 44 HAS, chlorophyll fluorescence decreased 56 and 36%, respectively, in methomyl-treated plants compared to controls (Fig. 2D). By 92 HAS, chlorophyll fluorescence increased nearly 25% in methomyl-treated plants compared to control plants.

## DISCUSSION

The different conditions in the two experiments (e.g. cultivars, soil types, planting dates, and spraying dates) were used to test whether responses to methomyl treatment were similar under differing conditions. The two experiments did respond similarly, although some minor differences occurred.

Photosynthetic injury in methomyl-treated plants was transient. Carbon exchange rates, stomatal conductance, and chlorophyll fluorescence decreased with methomyl treatment, but recovered (i.e. did not differ significantly from controls) by 100 HAS. Rubisco activity showed the same trend in experiment I (the 100 HAS readings of the experiment II were missing). The CER showed different trends in the two experiments. The lowest CER occurred at 28 HAS in experiment I, but did not occur till 78 HAS in experiment II. In both cases, the CER of methomyl-treated and control plants were not significantly different by 100 HAS. It remains unclear whether the different CER responses were due to cultivar or growing conditions, which may be an interesting question for further research.

A comparison of CER with other processes is instructive. Chlorophyll fluorescence recovered from methomyl treatment before CER did. In addition, chlorophyll fluorescence at 100 HAS was actually greater (numerically in experiment I, but significantly in experiment II) in methomyl-treated plants than in control plants, but CER of methomyl-treated plants never increased above those of controls. Because CER was measured on whole leaves, whereas chlorophyll fluorescence was measured on non-injured (i.e. not red or chlorotic) portions of leaves, the slower recovery of CER may simply reflect the inclusion of severely injured leaf area in the measurement. However, in experiment II, CER were still declining when chlorophyll fluorescence was increasing. Hence, chlorophyll fluorescence appears insufficient to explain completely the changes in CER.

Rubisco activity, like chlorophyll fluorescence, tended to recover before CER did. Also, the decreases in rubisco activity were never as severe as the decreases in CER. Furthermore, CER had already declined significantly at 4 HAS in both experiments; at that time, however, rubisco activity differed little between methomyl-treated and control plants. Thus, rubisco activity cannot wholly account for decreased CER.

Stomatal conductance and CER showed quite similar patterns, especially for the methomyl-treated plants. The relationship between stomatal conductance and CER was weaker in controls than in methomyl-treated plants, especially in experiment I. Apparently, in the unstressed controls, stomatal conductance was not the limiting factor in photosynthesis (at least in experiment I). The decreases in stomatal conductance were equal to or greater than the decreases in CER. Thus, stomatal conductance is a good candidate for explaining changes in CER. This conclusion is at variance with the findings of Wood and Payne (1986). In their study, various insecticides (including methomyl) decreased CER of pecan leaves, but generally did not affect stomatal conductance. Haile et al. (1999) also reported that besides a transient increase in photosynthesis in one year, there was no significant effects of the tested insecticides on other physiological parameters of either alfalfa *(Medicago sativa L.)* or soybean *[Glycine max (L.)]*. No significant methomyl effect was detected on lettuce *(Lactuca sativa L.)* either (Haile et al., 2000). Thus, plant species or even varieties may play a key role in how the changes in stomatal conductance would affect CER.

## CONCLUSIONS

Methomyl-induced injury to CER and related processes in cotton is transient, and the recovery generally occurring by 100 HAS. Further studies should examine effects of different cultivars and growing conditions on degree of injury and recovery rate.

All physiological processes studied showed decreases, followed by recovery. Although the three parameters may all affect CER, the changes in CER of cotton plants appear most closely related to changes in stomatal conductance.

